## Supplementary Tables for "Transcranial focused ultrasound-mediated neurochemical and functional connectivity changes in deep cortical regions in humans"

**Supplementary Table 1.** Neuronavigation parameters used and output of acoustic simulations of the dorsal anterior cingulate cortex TUS target.

| | Neuronavigation parameters | | | Simulated output | | | | | | | | | | |
|---|---|---|---|---|---|---|---|---|---|---|---|---|---|---|
| **ID** | **Transducer base coordinates** | **Target depth [mm]** | **Target coordinates** | **PPW** | **CFL** | **Maximum pressure coordinates** | **Focal depth [mm]** | **Maximum pressure [MPa]** | **MI** | **ISPPA [$W/cm^2$]** | **ISPTA [$mW/cm^2$]** | **Focal volume [$mm^3$]** | **Volume overlapping MRS voxel [$mm^3$]** | **Distance to COG of MRS voxel [mm]** |
| TS01 | 59,147,230 | 62.8 | 78,129,163 | 3 | 0.20 | 79,131,168 | 58.96 | 0.852 | 1.21 | 24.22 | 2422.38 | 553 | 371 | 5.83 |
| TS02 | 89,192,261 | 59.7 | 98,160,201 | 3 | 0.20 | 100,161,203 | 58.86 | 0.833 | 1.18 | 23.12 | 2311.90 | 500 | 351 | 2.47 |
| TS03 | 71,187,219 | 61 | 82,160,155 | 3 | 0.20 | 84,161,159 | 58.78 | 0.854 | 1.21 | 24.33 | 2433.03 | 442 | 333 | 3.84 |
| TS04 | 64,209,207 | 58.7 | 82,158,167 | 3 | 0.20 | 84,160,169 | 57.20 | 0.737 | 1.04 | 18.12 | 1812.12 | 647 | 455 | 7.44 |
| TS05 | 61,202,214 | 66.2 | 83,162,155 | 3 | 0.20 | 85,165,158 | 64.26 | 0.832 | 1.18 | 23.07 | 2307.32 | 694 | 373 | 11.40 |
| TS06 | 36,128,235 | 68 | 83,125,175 | 3 | 0.20 | 82,126,180 | 63.90 | 0.768 | 1.09 | 19.66 | 1966.05 | 802 | 537 | 4.38 |
| TS07 | 67,182,240 | 61.7 | 98,173,177 | 3 | 0.20 | 97,175,183 | 56.34 | 0.860 | 1.22 | 24.67 | 2466.57 | 422 | 348 | 5.84 |
| TS08 | 90,144,236 | 60.1 | 93,137,167 | 3 | 0.20 | 94,138,168 | 60.42 | 0.864 | 1.22 | 24.90 | 2490.25 | 418 | 277 | 8.88 |
| TS09 | 50,159,233 | 62.1 | 80,143,169 | 3 | 0.20 | 81,142,169 | 58.22 | 0.810 | 1.15 | 21.89 | 2189.30 | 661 | 464 | 3.15 |
| TS10 | 47,192,219 | 67.1 | 81,155,164 | 3 | 0.20 | 86,154,161 | 72.29 | 0.692 | 0.98 | 15.96 | 1596.28 | 1122 | 505 | 12.38 |
| TS11 | 70,161,234 | 59.6 | 83,146,168 | 3 | 0.20 | 85,147,170 | 59.16 | 0.877 | 1.24 | 25.61 | 2560.78 | 413 | 282 | 5.79 |
| TS12 | 88,179,267 | 61.7 | 98,157,200 | 3 | 0.20 | 100,159,203 | 69.19 | 0.918 | 1.30 | 28.08 | 2807.87 | 456 | 330 | 3.86 |
| TS13 | 73,162,232 | 55.1 | 80,140,170 | 3 | 0.20 | 82,142,173 | 64.07 | 0.847 | 1.20 | 23.89 | 2389.25 | 396 | 291 | 3.89 |
| TS14 | 102,160,268 | 57.5 | 99,152,201 | 3 | 0.20 | 100,153,203 | 66.55 | 0.846 | 1.20 | 23.84 | 2383.75 | 422 | 302 | 4.60 |
| TS16 | 77,152,256 | 55.1 | 85,139,193 | 3 | 0.20 | 86,141,195 | 52.68 | 0.873 | 1.23 | 25.39 | 2538.83 | 300 | 249 | 3.14 |
| TS17 | 57,150,242 | 58.7 | 84,141,181 | 3 | 0.20 | 86,143,180 | 60.40 | 0.748 | 1.06 | 18.64 | 1863.99 | 638 | 350 | 8.75 |
| TS18 | 66,127,255 | 64.6 | 85,125,187 | 3 | 0.20 | 87,125,186 | 63.86 | 0.800 | 1.13 | 21.33 | 2132.55 | 697 | 350 | 9.83 |
| TS19 | 50,168,245 | 63.1 | 83,144,185 | 3 | 0.20 | 84,146,188 | 61.58 | 0.805 | 1.14 | 21.61 | 2160.75 | 705 | 443 | 6.19 |
| TS20 | 62,172,228 | 53.9 | 84,147,173 | 3 | 0.20 | 86,148,176 | 52.95 | 0.724 | 1.02 | 17.45 | 1745.15 | 550 | 397 | 2.21 |
| TS21 | 69,150,244 | 60.1 | 83,140,177 | 3 | 0.20 | 84,141,180 | 58.24 | 0.817 | 1.16 | 22.26 | 2226.19 | 452 | 338 | 2.81 |
| TS23 | 59,166,219 | 53.5 | 83,147,160 | 3 | 0.20 | 84,149,166 | 51.77 | 0.809 | 1.14 | 21.79 | 2179.03 | 354 | 303 | 3.13 |
| TS24 | 73,158,229 | 52.1 | 85,145,169 | 3 | 0.20 | 86,147,172 | 50.43 | 0.903 | 1.28 | 27.21 | 2720.60 | 260 | 230 | 2.68 |
| TS25 | 77,156,218 | 60.4 | 83,147,150 | 3 | 0.20 | 84,149,155 | 55.78 | 0.820 | 1.16 | 22.40 | 2240.25 | 449 | 295 | 6.17 |
| TS26 | 70,177,222 | 56.3 | 83,145,165 | 3 | 0.20 | 85,147,170 | 52.98 | 0.846 | 1.20 | 23.88 | 2387.83 | 387 | 314 | 2.91 |

**Supplementary Table 2.** Neuronavigation parameters used and output of acoustic simulations of the posterior cingulate cortex TUS target.

| | Neuronavigation parameters | | | Simulated output | | | | | | | | | | |
|---|---|---|---|---|---|---|---|---|---|---|---|---|---|---|
| **ID** | **Transducer base coordinates** | **Target depth [mm]** | **Target coordinates** | **PPW** | **CFL** | **Maximum pressure coordinates** | **Focal depth [mm]** | **Maximum pressure [MPa]** | **MI** | **ISPPA [W/cm$^2$]** | **ISPTA [mW/cm$^2$]** | **Focal volume [mm$^3$]** | **Volume overlapping MRS voxel [mm$^3$]** | **Distance to COG of MRS voxel [mm]** |
| TS01 | 66,3,201 | 68 | 79,67,159 | 3 | 0.20 | 79,66,161 | 67.29 | 0.793 | 1.12 | 20.97 | 2097.48 | 743 | 457 | 6.55 |
| TS02 | 106,57,262 | 75.1 | 98,100,194 | 3 | 0.20 | 99,101,196 | 74.01 | 0.840 | 1.19 | 23.52 | 2351.98 | 867 | 457 | 6.65 |
| TS03 | 75,111,231 | 75.8 | 82,104,150 | 3 | 0.20 | 82,106,153 | 73.45 | 0.856 | 1.21 | 24.44 | 2443.96 | 720 | 407 | 10.08 |
| TS04 | 41,67,222 | 72.2 | 83,94,160 | 3 | 0.20 | 81,94,165 | 67.57 | 0.869 | 1.23 | 25.15 | 2515.45 | 626 | 476 | 2.88 |
| TS05 | 74,83,244 | 69.6 | 81,105,170 | 3 | 0.20 | 82,109,167 | 74.21 | 0.743 | 1.05 | 18.42 | 1842.31 | 986 | 387 | 11.59 |
| TS06 | 60,21,215 | 69.4 | 82,71,160 | 3 | 0.20 | 82,73,161 | 69.89 | 0.821 | 1.16 | 22.46 | 2246.21 | 741 | 454 | 4.79 |
| TS07 | 89,79,234 | 76.6 | 99,113,160 | 3 | 0.20 | 100,113,162 | 74.73 | 0.839 | 1.19 | 23.47 | 2346.53 | 839 | 458 | 6.01 |
| TS08 | 60,46,215 | 72 | 83,90,153 | 3 | 0.20 | 85,95,151 | 77.29 | 0.761 | 1.08 | 19.32 | 1932.00 | 990 | 374 | 14.48 |
| TS09 | 91,42,220 | 73.3 | 84,89,155 | 3 | 0.20 | 85,89,157 | 72.13 | 0.775 | 1.10 | 20.02 | 2002.43 | 998 | 490 | 3.81 |
| TS10 | 80,118,255 | 82 | 84,107,174 | 3 | 0.20 | 86,110,173 | 80.64 | 0.795 | 1.12 | 21.04 | 2104.16 | 1031 | 526 | 6.91 |
| TS11 | 87,52,213 | 61.7 | 84,97,158 | 3 | 0.20 | 85,97,161 | 59.95 | 0.877 | 1.24 | 25.62 | 2561.52 | 481 | 311 | 7.15 |
| TS12 | 93,49,254 | 64 | 98,94,197 | 3 | 0.20 | 99,95,197 | 64.58 | 0.840 | 1.19 | 23.49 | 2349.12 | 604 | 362 | 5.29 |
| TS13 | 68,50,237 | 68.5 | 79,81,167 | 3 | 0.20 | 79,83,168 | 69.61 | 0.813 | 1.15 | 22.06 | 2205.56 | 991 | 577 | 6.81 |
| TS14 | 104,64,256 | 64 | 99,100,190 | 3 | 0.20 | 100,100,193 | 64.12 | 0.852 | 1.20 | 24.20 | 2420.12 | 587 | 382 | 4.19 |
| TS16 | 50,51,225 | 71 | 82,82,171 | 3 | 0.20 | 83,85,168 | 67.01 | 0.947 | 1.34 | 29.90 | 2989.81 | 494 | 219 | 11.92 |
| TS17 | 77,52,244 | 68.2 | 83,85,175 | 3 | 0.20 | 84,89,173 | 72.72 | 0.758 | 1.07 | 19.14 | 1914.23 | 833 | 439 | 7.29 |
| TS18 | 72,9,207 | 75.8 | 85,78,164 | 3 | 0.20 | 85,81,165 | 77.87 | 0.838 | 1.19 | 23.41 | 2341.16 | 860 | 496 | 8.43 |
| TS19 | 30,51,226 | 75.5 | 84,83,174 | 3 | 0.20 | 87,87,171 | 80.59 | 0.777 | 1.10 | 20.14 | 2013.79 | 967 | 538 | 7.69 |
| TS20 | 69,60,235 | 62.7 | 82,90,171 | 3 | 0.20 | 83,92,172 | 63.29 | 0.785 | 1.11 | 20.53 | 2052.77 | 607 | 390 | 3.31 |
| TS21 | 65,34,218 | 68.2 | 82,82,162 | 3 | 0.20 | 83,83,165 | 66.22 | 0.774 | 1.09 | 19.99 | 1998.61 | 747 | 455 | 3.02 |
| TS23 | 65,67,219 | 56.1 | 83,92,160 | 3 | 0.20 | 84,94,161 | 56.94 | 0.868 | 1.23 | 25.11 | 2511.04 | 369 | 303 | 2.54 |
| TS24 | 86,29,196 | 64.3 | 84,87,151 | 3 | 0.20 | 85,84,155 | 59.44 | 0.873 | 1.24 | 25.43 | 2542.69 | 497 | 356 | 3.90 |
| TS25 | 77,47,198 | 63.2 | 84,92,141 | 3 | 0.20 | 84,92,144 | 61.69 | 0.868 | 1.23 | 25.08 | 2508.37 | 477 | 350 | 2.59 |
| TS26 | 66,58,227 | 69.9 | 84,91,159 | 3 | 0.20 | 84,93,160 | 69.90 | 0.851 | 1.20 | 24.16 | 2416.44 | 642 | 415 | 4.38 |
