## Supplementary Figure 1 for "Transcranial focused ultrasound-mediated neurochemical and functional connectivity changes in deep cortical regions in humans"

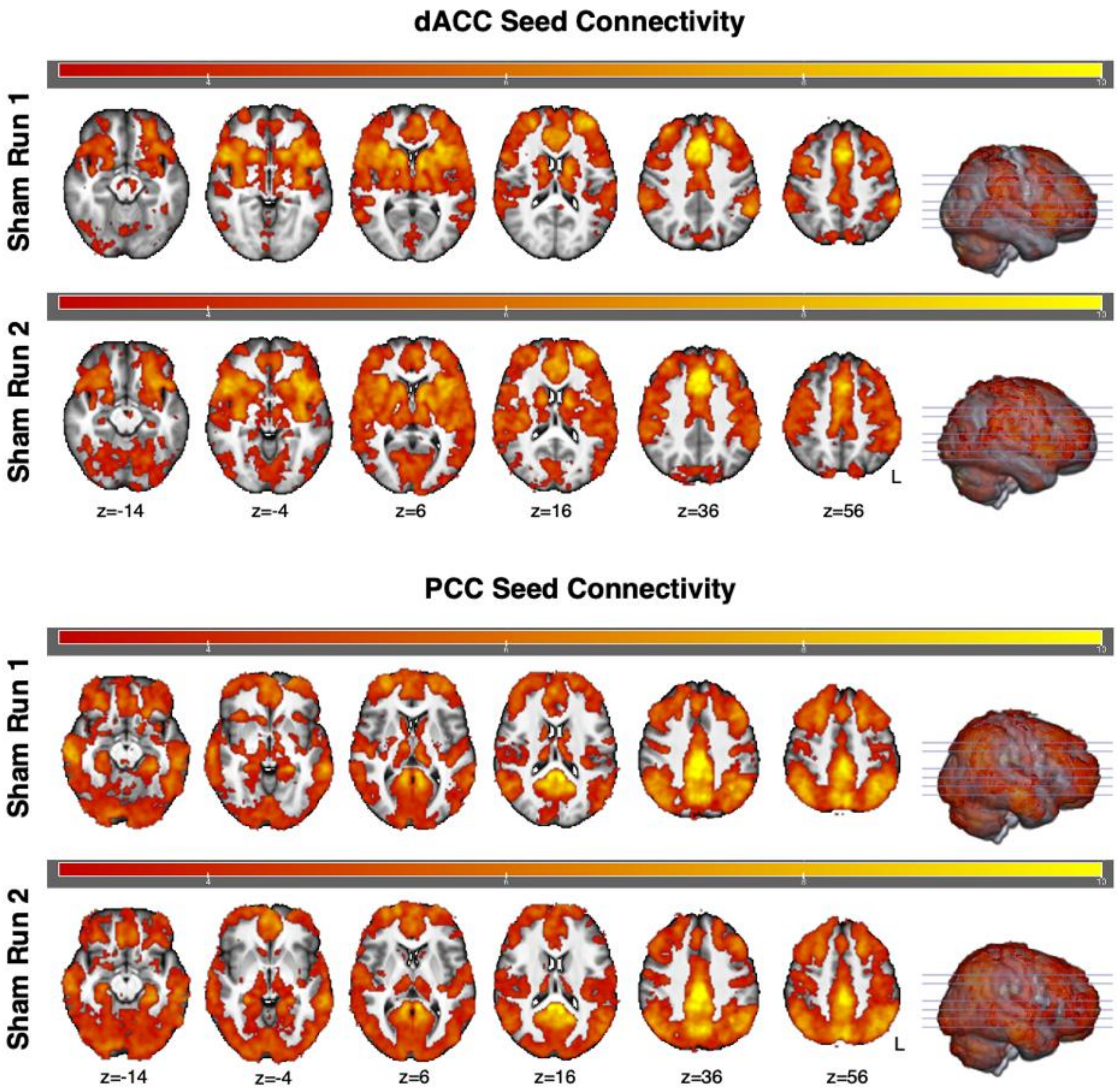


**Supplementary Figure 1.** Seed-based functional connectivity of the dACC (left) and PCC (right) in the early rsfMRI run during the sham session, Sham Run 1, at approximately 13 minutes post-TUS, and the late rsfMRI run during the sham session, Sham Run 2, at approximately 46 minutes post-TUS. Whole-brain maps are overlaid on the average T1-weighted MRI of all participants. There were no significant differences between the early and late rsfMRI runs during the sham session for either the dACC or the PCC seed.
