## Supplementary Figure 2 for "Transcranial focused ultrasound-mediated neurochemical and functional connectivity changes in deep cortical regions in humans"

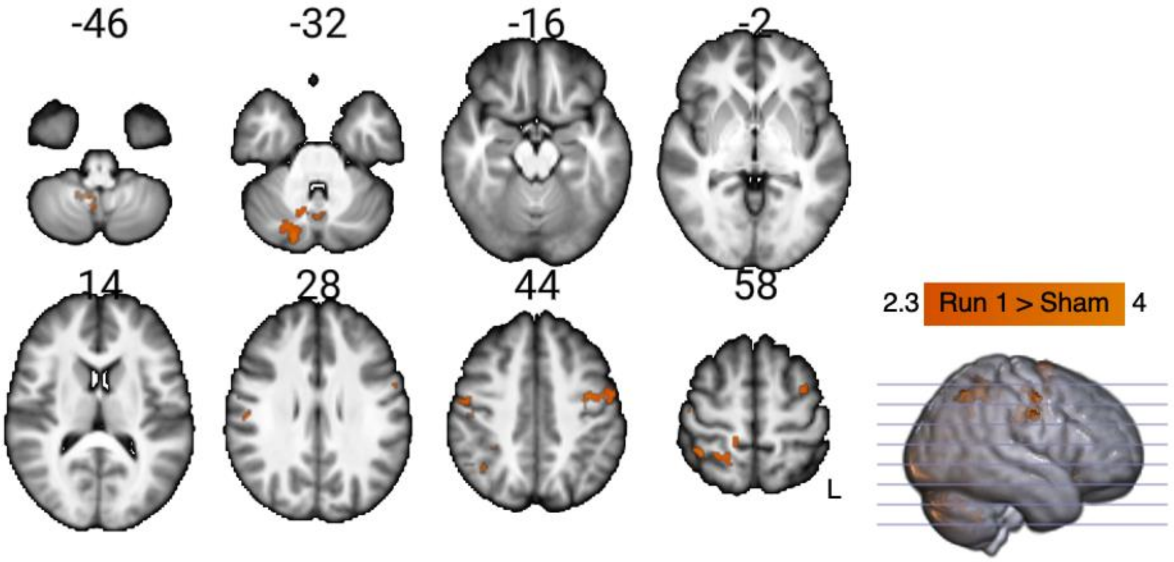


**Supplementary Figure 2.** Functional connectivity changes with the dACC seed after TUS was applied to the PCC during the early rsfMRI run (i.e., approximately 13 minutes post-TUS) compared with the average of the two rsfMRI runs during the sham session. Whole-brain maps illustrate regions showing increased functional connectivity (Z statistics; cluster-corrected at $p < 0.05$).
