## Supplementary Figure 3 for "Transcranial focused ultrasound-mediated neurochemical and functional connectivity changes in deep cortical regions in humans"

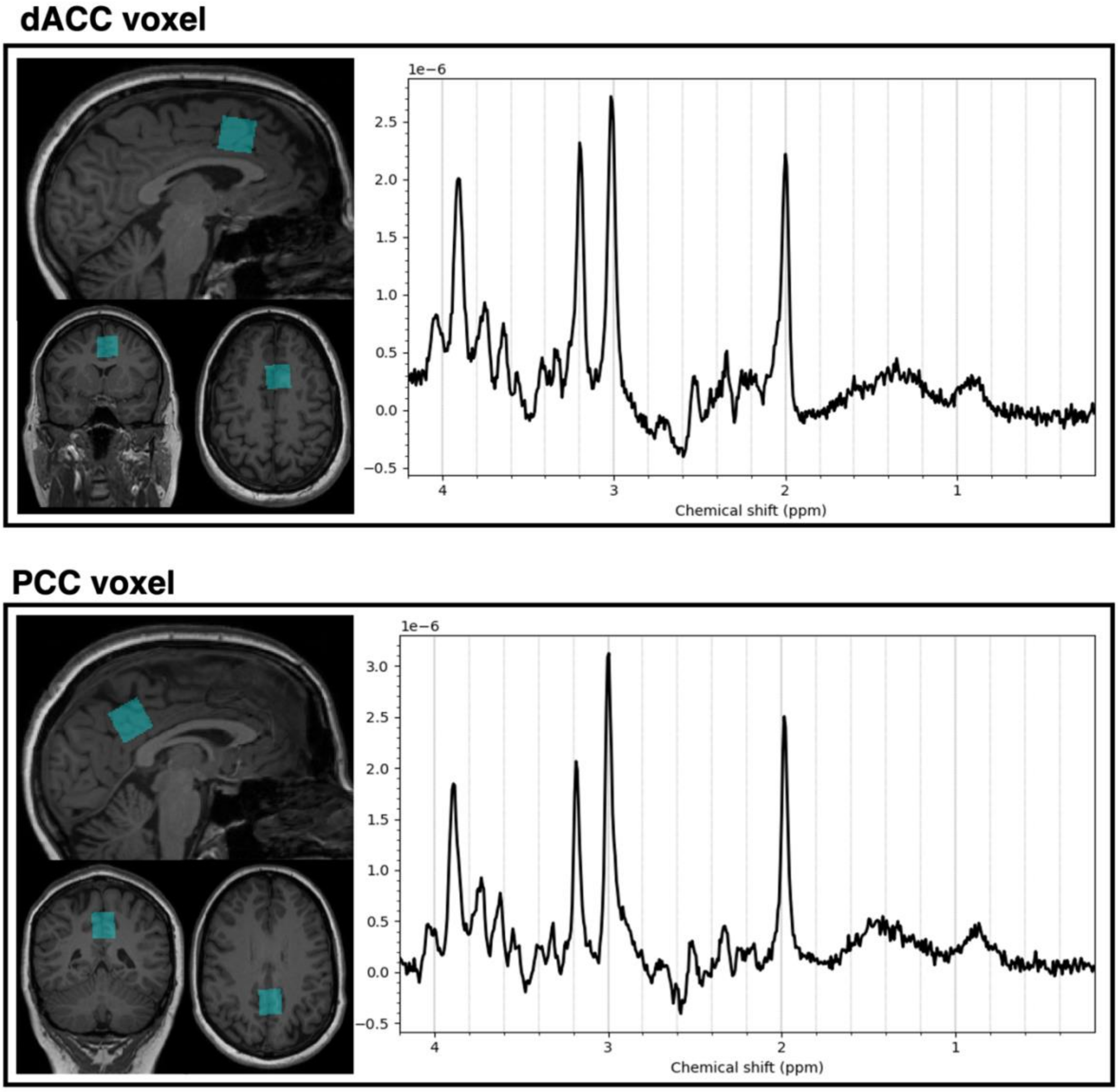


**Supplementary Figure 3.** Representative spectrum and voxel location acquired from the dorsal anterior cingulate cortex (dACC; left) and the posterior cingulate cortex (PCC; right) in one individual. The chemical shift axis is labeled in ppm units.
